# Are human connectomes heritable?

**DOI:** 10.1101/2023.04.02.532875

**Authors:** Jaewon Chung, Eric W. Bridgeford, Michael Powell, Derek Pisner, Ting Xu, Joshua T. Vogelstein

## Abstract

A complete understanding of human behavior and disease depends upon our ability to parse genetic and environmental influences in the human brain. The heritability of a trait quantifies the degree of its variability due to genetic influences. Classical approach for quantifying heritability operate on simple traits, and sometimes do not properly control for other potential sources of variation, such as age or sex. We therefore develop Causal Heritability of Networks (CHaiN) to rigorously quantify heritability of human brain networks (i.e., connectomes). We applied CHaiN to 1024 anatomical connectomes derived from the Human Connectome Project. Connectomes appeared to be heritable, but heritability was insignificant once we addressed variability within networks. These results suggest that previous conclusions on connectome heritability may be driven by the shared network structures, and highlights the importance of modeling networks and other sources of variability when studying heritability of connectomes.

## 1 Introduction

Many common human traits and diseases cluster in families, and are believed to be influenced by several genetic and environmental effects. Heritability quantifies the total variability in a given trait or disease and is attributable to genetic factors and not environmental or stochastic events [1]. Thus, heritability is a key factor in predicting disease risk from an individual’s history [2], setting bounds on the ability of genetics to predict disease [2], providing justifications for further genetic studies, and developing genetic treatments for heritable diseases, such as sickle cell anemia [3–7].

Numerous methods of estimating heritability, that is, partitioning phenotypic variance into genetic and environmental components, have been applied to various “simple” traits in twin studies. The intraclass correlation (ICC) [8], which estimates the amount of variation that is common to the group as a proportion of variation in the population, and Falconer’s method [9], which is the difference in ICC in identical and fraternal twins of a given trait, have been used to show significant heritability in human personalities [10–12], anatomy [13–16], and intelligence [17, 18]. Structural equation models (SEMs) have also been used to estimate the heritability of phenotypically complex diseases such as schizophrenia and Alzheimer’s [19, 20]. SEMs, in contrast to ICC, can incorporate multivariate inputs, account for covariates such as sex and age, incorporate familial relationships such as step siblings and parents, and provide more accurate estimates of heritability [21]. Although SEMs are extremely useful in human genetics [22], they exhibit several limitations. For example, phenotypic data are typically assumed to be normally distributed, which is often violated in complex traits, and phenotypic variance can only be modeled as a linear relationship among its variance components.

Recent advances in neuroimaging have enabled the investigation of heritability in complex neural traits, particularly brain connectivity [23–27]. Connectomes, which model functional and/or structural connections between brain regions, provide a powerful framework for representing brain connectivity topology through graph theory [28–30]. However, these earlier studies faced limitations that can hinder clear interpretations. For instance, conclusions about heritability between network metrics (e.g., average path length vs. clustering coefficient) are obscured by the inherent correlations between these measures [31]. Furthermore, the extensive array of graph-theoretical metrics available for connectome analysis raises statistical challenges. Selecting the most appropriate metrics to accurately assess brain connectivity heritability, while mitigating the risk of false discoveries from multiple comparisons, remains an open question. Though graph-theoretical features offer intuitive characterizations of individual connectomes, no single set can fully encapsulates network topology [31]. Finally, the non-Euclidean and non-Gaussian nature of connectome data motivates specialized analytical approaches. Traditional methods like SEMs, which rely on Euclidean and Gaussian assumptions, are inappropriate. Violating these assumptions risks erroneous inferences and compromises scientific reproducibility.

In this work, we frame the question of human structural connectome heritability as a causal inference problem: do genetic changes directly influence connectome variations? To address this, we introduce Causal Heritability of Connectomes (CHaiN), which defines a causal model and proposes a novel method using distance correlations to detect and estimate this heritability. This approach accommodates both linear and non-linear dependencies between genetics and connectomes (Figure 1). Furthermore, we offer three models for connectome comparison, each with distinct assumptions that reveal different potential aspects of heritability. These models systematically remove common network structures with increasing complexity, allowing us to isolate heritable effects beyond these shared patterns.

**Figure 1.**
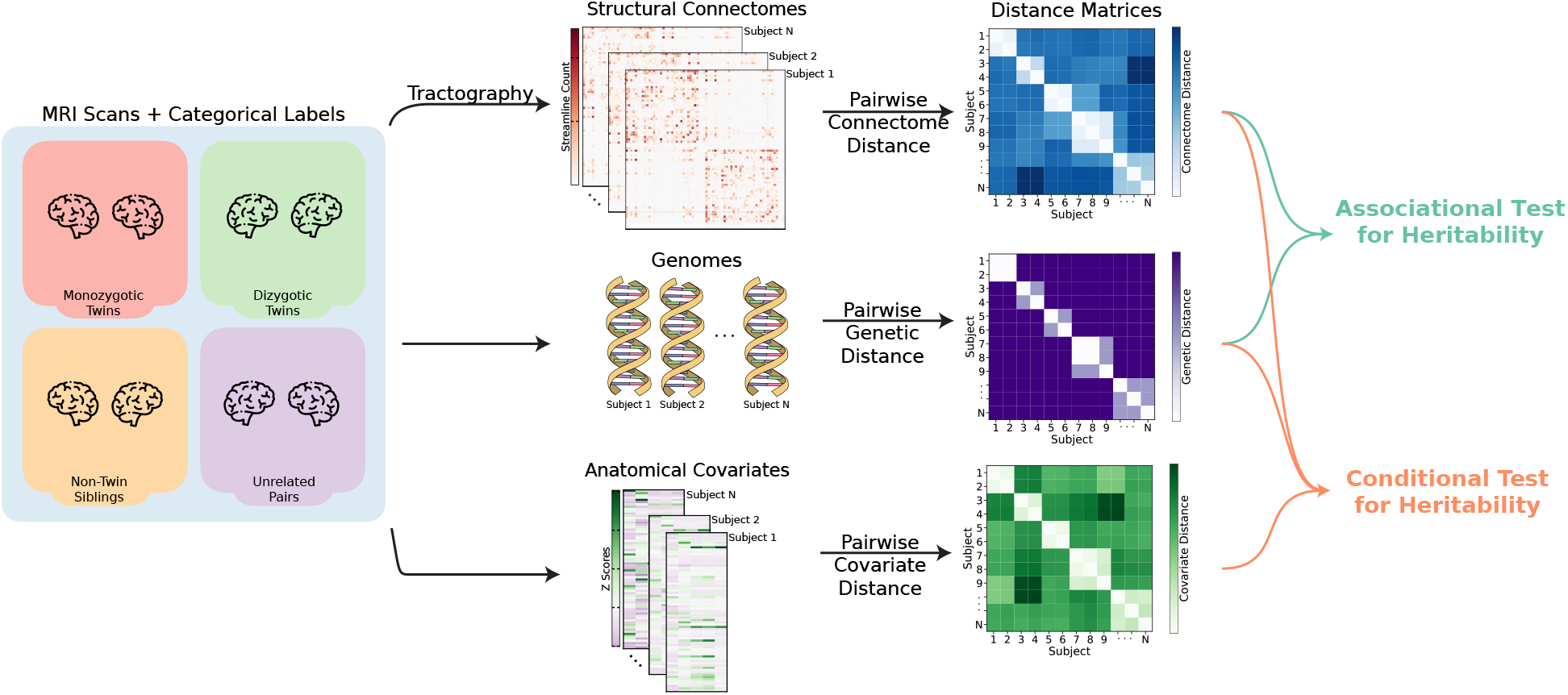
Overview of the framework for measuring heritability of connectomes. (*Left*) Diffusion and structural magnetic resonance images (dMRI and sMRI) are processed to generate connectomes. Connectomes are defined on a common set of vertices using brain parcellations. Each pair of individuals has a label corresponding to whether the pair is a monozygotic twin, dizygotic twin, non-twin siblings, or unrelated (e.g. not sharing both mother and father). (*Center left*) For each subject, connectomes and covariates are estimated from MRI scans and phenotypic data. (*Center right*) For each modality (e.g. connectomes, genomes, and covariates), distances are computed between all possible pairs of subjects using the corresponding distance function. This results in distance matrices that are input to subsequent hypothesis tests. (*Right*) Associational effect for connectomic heritability is measured using distance correlation (Dcorr). Conditional effect for connectomic heritability is measured using conditional distance correlation (CDcorr). Significance tests provide evidence to reject the null that genomes and connectomes are not related.

Using Human Connectome Project data, we demonstrate that structural connectomes are heritable only up to some connectome model commonalities, even when rigorously controlling for potential confounders (e.g., age, neuroanatomy, sex). Our methodology lays the foundation for future research on how genetic factors influence complex brain phenotypes. Crucially, it highlights the importance of modeling the inherent structure of connectomes and the need for specialized analytical methods that respect the non-Euclidean and non-Gaussian nature of brain connectivity data.

## 2 Causal Heritability of Connectomes (CHaiN)

### 2.1 Model of Heritability in Twin Datasets

We focus on demonstrating the presence of heritability for structural connectomes in a twin neuroimaging study (see Figure 1 for an overview). Figure 2 illustrates a directed acyclic graph (DAG) representing a twin study, where measurements are obtained from individuals across multiple families. In the DAG, the directionality of arrows (e.g. *Z → Y*) indicates a potential causal influence of variable *Z* on variable *Y*. Ideally, we aim to estimate the causal effect of genetic variation on structural connectome variation while controlling for all possible covariates, observed or unobserved. However, a key limitation is that estimating this pure causal effect is infeasible in practice, as it requires knowledge of both measured and unmeasured confounders (the causal framework is described in more detail in Section 5.1).

**Figure 2.**
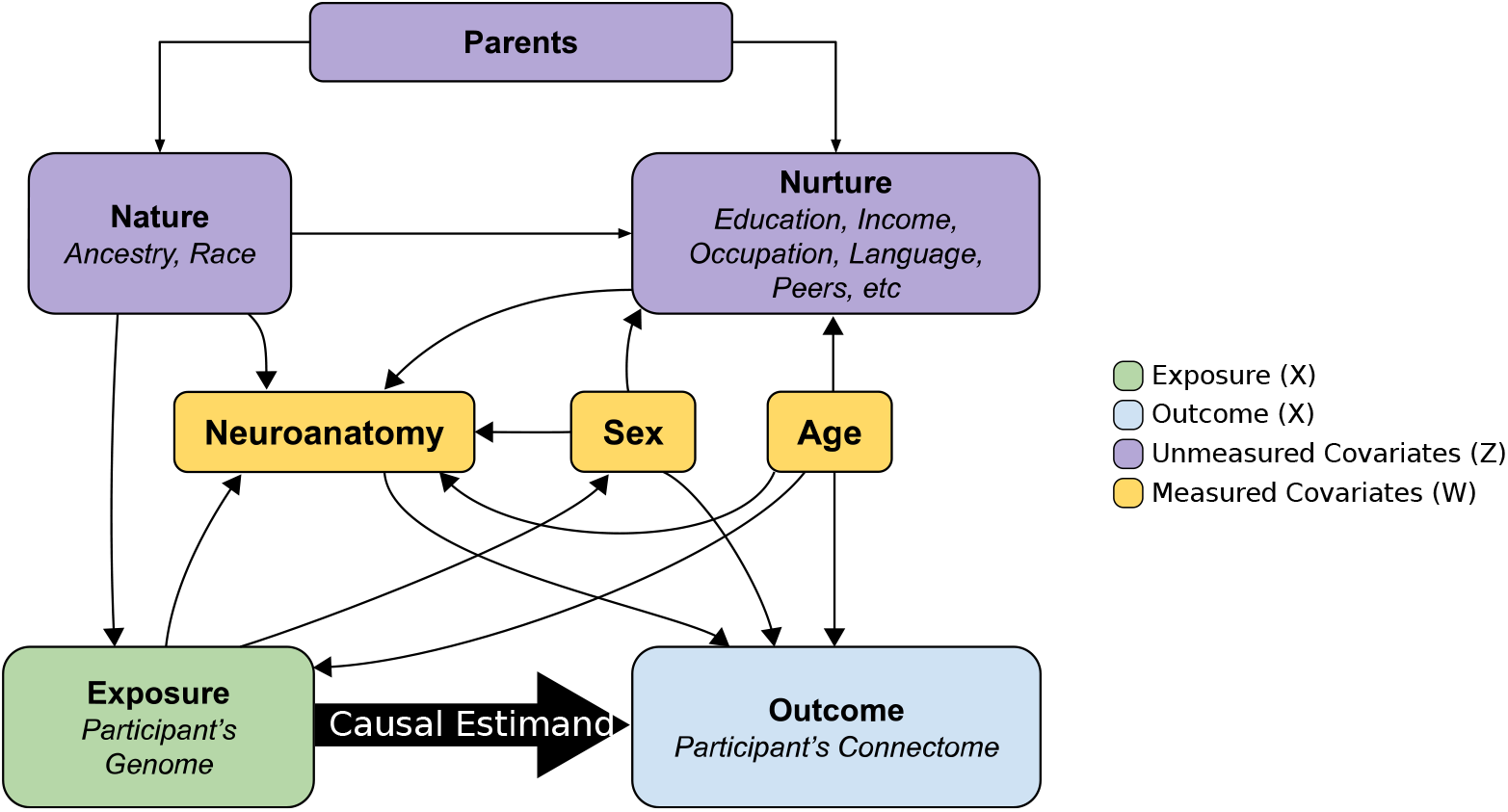
Causal Graph of Study Covariates. Causal directed acyclic graph (DAG) presents a visual representation of the potential relationships between the genome and connectome, in the context of a twin study. The descriptions depict the various attributes that may be considered as exposures, outcomes, and covariates in such a study. In the absence of confounding, associational effects can be considered as causal effects. However, conditional effects can only be considered as causal when the covariates measured are sufficient to close all backdoor paths.

Given the data available from twin neuroimaging studies, what models can help us provide evidence of heritability?

#### 1 Associational Heritability

The simplest model examines whether the genome is associated with connectomes. However, in real studies, both genome and connectomes are often influenced by other covariates. This makes estimated associational effects potentially unreliable, as they may conflate true genetic effects with those of correlated factors. For instance, if brain size relates to both genetics and structural connectomes, we cannot easily distinguish whether observed associations are due to genetics or neuroanatomy.

#### 2 Conditional Heritability

This model addresses the limitations of associational heritability. It asks whether differences in the outcome (structural connectomes) are associated with differences in genetic exposure, after conditioning on measured covariates. Observing such differences strengthens the argument for causal genetic influence.

Conditional heritability can be considered equivalent to **causal heritability** under a sufficient condition: the strong ignorability assumption [32]. In twin studies, a simplifying assumption often made is the “equal environment assumption”, that monozygotic (MZ) and dizygotic (DZ) twins share environmental factors to a similar degree by virtue of their study design. Additionally, in non-twin siblings, some overlap in covariate distributions is likely (e.g., if siblings share a household). Assuming that other experimental design considerations ensure overlap in covariates such as age, we can posit that neuroanatomy, sex, and age close all possible non-causal pathways that can create spurious correlations (e.g. close potential backdoor paths; see Section 5.1 for details). If these conditions hold, the conditional heritability in our study can be interpreted as causal. Next, we describe a twin neuroimaging study that will be used for subsequent analysis.

### 2.2 Twin Dataset for Discovering Connectomic Heritability

Human Connectome Project (HCP) Young Adult study, acquired by Washington University in St. Louis (WUSTL) and the University of Minnesota (Minn), provides high-quality open-access neuroimaging data from standardized protocols that minimize potential batch effects [33, 34]. Moreover, the inclusion/exclusion criterion guarantees that individuals with specific factors, like neurological diseases, which could potentially impact brain connectivity, are not included in the study. We used human brain diffusion (dMRI) and structural magnetic resonance imaging (sMRI) data to estimate structural connectomes (see 5.2 for more details on network construction). Out of the 1206 participants, a total of 1024 participant brain images were processed. Families left with only one subject were discarded, and half-siblings were discarded. The data set includes 322 (196 females; ages 22 *−* 36 years old, mean = 29.6 and SD = 3.3) monozygotic twins, 212 (125 females; ages 22 *−* 36 years old, mean = 28.6 and SD = 3.4) dizygotic twins, and 490 (237 females; ages 22 *−* 37 years old, mean = 28.3 and SD = 3.9) non-twin siblings. Figure 3 shows visualizations of the average connectome and the difference in two most different subjects for the Desikan parcellation.

**Figure 3.**
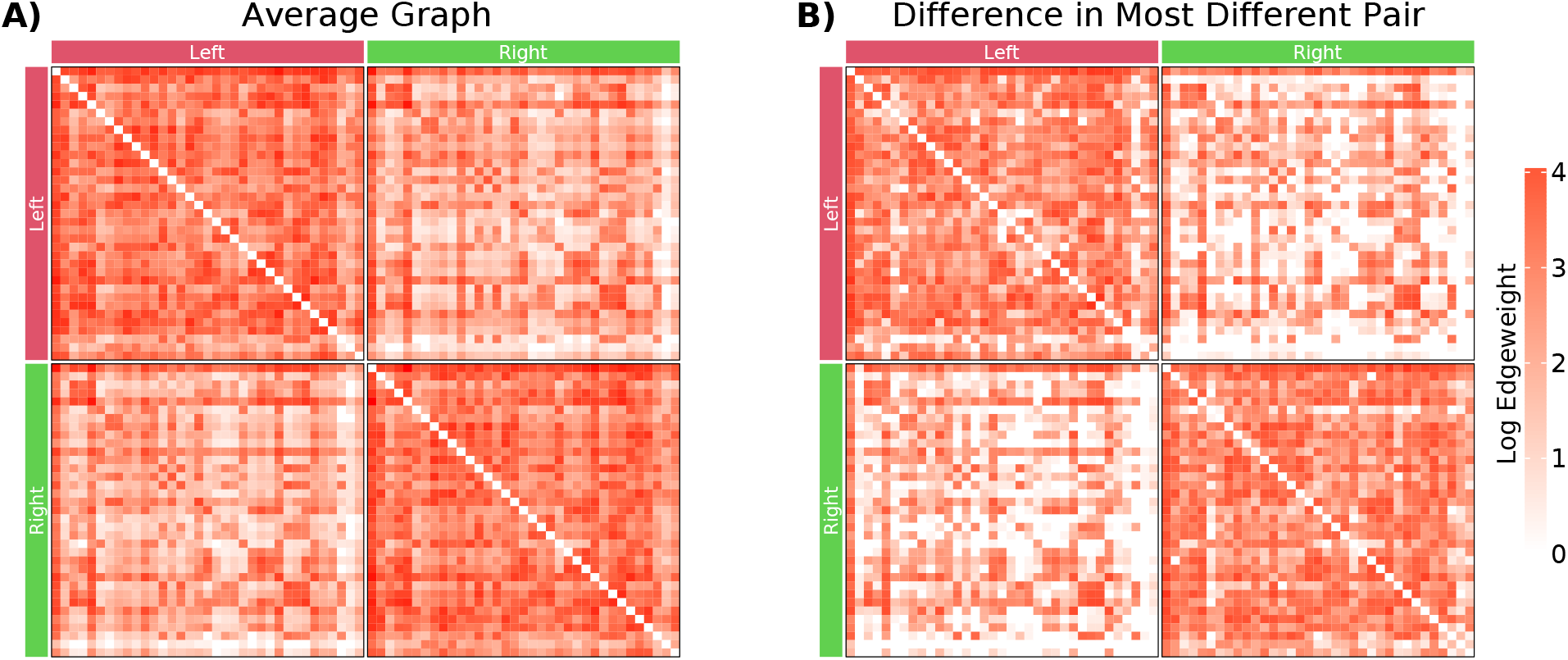
Visualization of connectomes as adjacency matrices using the projected Desikan parcellation with hemispheric labels. The projected Deskian parcellation was chosen for visualization purposes due to the low number of vertices (*n* = 70). **(A)** Average connectome of all subjects with log-transformed edge weights. **(B)** Absolute difference of connectomes from the most different pair of subjects with log-transformed edge weights.

Next, we detail various ways of measuring differences between pairs of connectomes and what aspects of connectomes they potentially characterize.

### 2.3 Methods for Comparing Connectomes

Connectomes have inherent structures and dependencies that complicate their comparison, along with noise present in the observed data. To address this, we introduce the concept of “latent positions” – a set of estimated variables that capture the underlying topologies and dependencies within a connectome. By comparing latent positions across connectomes, we gain a more nuanced understanding of their differences (refer to Section 5.4 for further information on the network model and techniques for estimating these latent positions). This approach allows us to move beyond simple structural comparisons, revealing deeper distinctions within the networks themselves. Given these representations, we propose three different models for computing distances between connectomes (see Section 5.5 for more mathematical details):

#### 1 Exact differences

This method measures all differences between latent positions, with differences in the latent positions implying differences in the connectomes themselves. Consequently, a hypothesis test based on an exact methods aims to test whether there is a correlation between the actual differences in connectomes and the differences in the genomes.

#### 2 Global differences

This method examines whether the latent positions of one connectome are a scaled version of the other. For example, if the estimated number of edges in male connectomes are consistently larger than those in females, we have no way of differentiating whether significant findings from the exact model are a result of differences in scaling or differences in the fundamental structure of the connectomes themselves. Thus, hypothesis tests based on global difference test whether differences in connectomes after adjusting for scaling differences correlate with differences in genomes. The global model assumes that the scaling factor is the same across all regions, meaning that any differences in scaling are not region-specific.

#### 1 Vertex differences

This method is similar to the global differences, but it allows for each vertex to be scaled differently. The idea behind this approach is that some vertices may have a greater impact on the overall network than others, so scaling them differently can provide a more accurate representation of the network. Consider the examples of how brain regions connect with each other. Regions in the same hemisphere are more likely to be connected than across hemispheres. Even within the same hemisphere, different regions may have distinct preferences for forming connections with other specific regions. Thus, hypothesis tests based on the vertex differences aim to determine whether there are significant differences in connectomes after accounting for differences in vertex-wise scaling.

To demonstrate the utility of these methods for measuring differences, we simulated connectomes with various latent positions as shown in Figure 4 (see Section A for technical details of these simulations). Columns I and II display four pairs of connectomes, each with specific latent position models. Rows (A-C) share the same underlying latent positions, while row (D) has different latent positions. When the connectome model can account for the differences in latent positions between the subjects, we observe a “uniform” structure in the differences, with no visible pattern. For the exact model (column III), distances increase as we move down the rows because it cannot account for any scaling, whether global or vertex-wise. We observe patterns in the differences, where red indicates edges in which subject 1 tended to have more edges, while blue shows edges where subject 1 tended to have fewer edges. For instance, for each simulation, when subject 1 tended to have more edges (row B.I-II), we observe uniform red coloring in the differences (row B.III). Similarly, for the global model, we only see patterns in differences when the differences in latent positions are due to vertex scaling, as shown by the gradients of red and blue (C.IV and D.IV). The main takeaway is that any hypothesis tests based on these models can only make claims within the limits of their ability to explain the differences in the pairs of connectomes.

**Figure 4.**
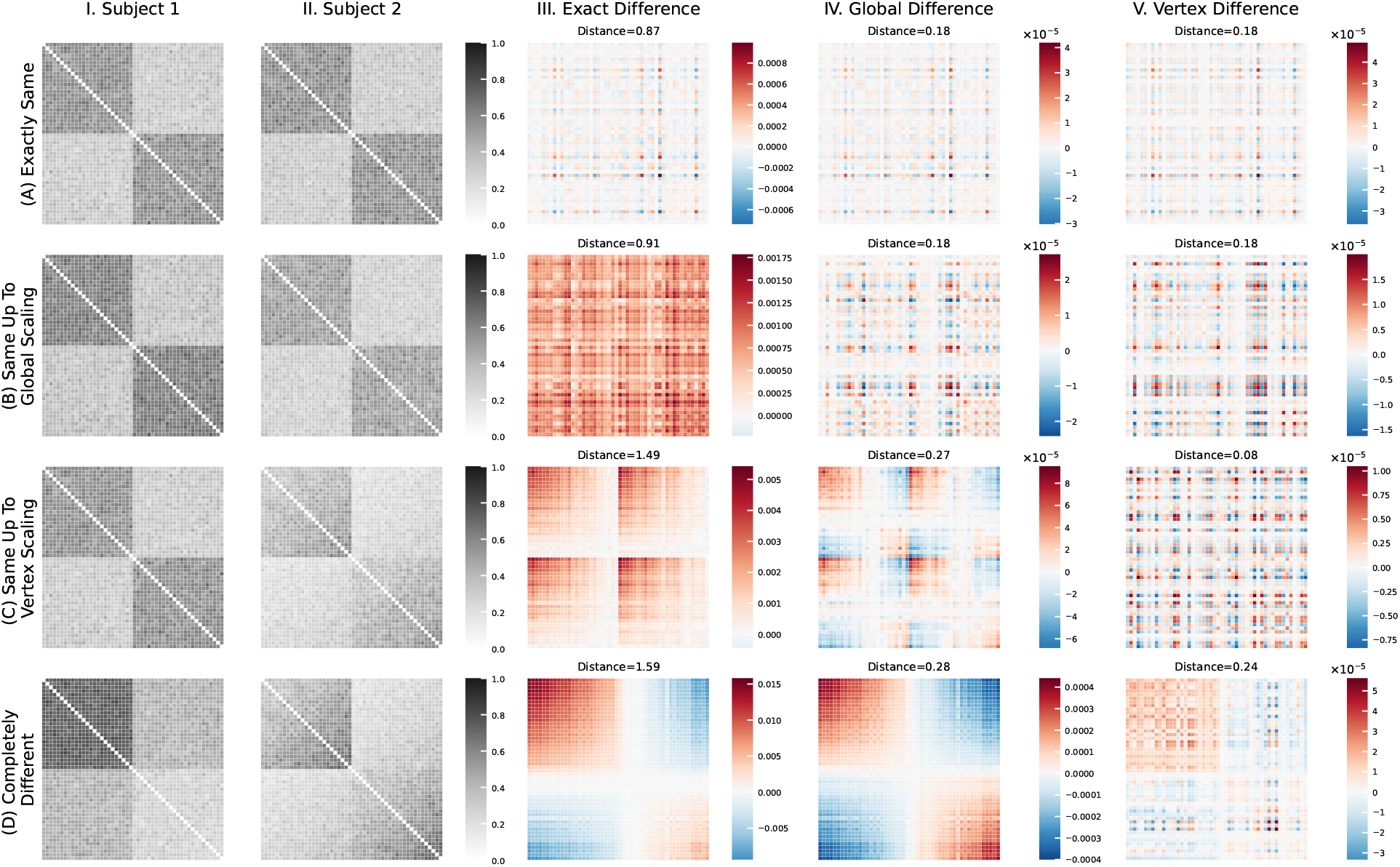
Simulations demonstrate the utility of connectome models. Connectomes with 50 vertices were simulated 100 times. Columns (I-II) display the average simulated connectomes, while columns (III-V) illustrate the differences in latent positions averaged over these simulations. In columns (III-V), red corresponds to when subject 1 had more edges than subject 2, while blue corresponds to when subject 1 had fewer edges. Rows (A-D) correspond to various parameters used to generate the connectomes. Connectomes in rows (A-C) share the same latent positions, subject to scaling transformations, whereas connectomes in row (D) have distinct latent positions, conditional on scaling transformations. When the connectome models accurately represent the generative process, the difference matrix displays a “uniform” structure without discernible patterns (A.III-V, B.IV-V, C.V). However, when connectome models are not appropriate, patterns emerge, as indicated by the increased distance values (B.III, C.III-VI, D.III-V).

## 3 Results

### 3.1 Associational Tests for Connectomic Heritability

We begin the examination of associational heritability by first comparing connectomes from monozygotic and same-sex dizygotic twins. By limiting the comparison to these two groups, we minimized the impact of environmental factors (e.g. Nurture in Figure 2) on the connectomes and assessed the ability of connectomes to capture variations in the genome. Shared and non-shared environmental influences may arise due to differences in age or sex, which can lead to differences in upbringing or educational experiences. By controlling for such factors, we could reasonably attribute any differences in connectomes to variations in genomes rather than environmental factors. Moreover, if the differences in connectomes between monozygotic twins were “smaller” than those between dizygotic twins, it would suggest that connectomes can indeed capture variations in genomes.

The distribution of connectome distances is shown in Figure 5A.I-III by the blue and yellow rows, revealing that the medians of monozygotic twins (*N* = 127) are smaller than those of same-sex dizygotic twins (*N* = 75) for all three connectome models. To determine the significance of this difference, we perform a test for an associational effect using Dcorr, yielding *p <* 10^*−*3^ for all models (Figure 5B.I).

**Figure 5.**
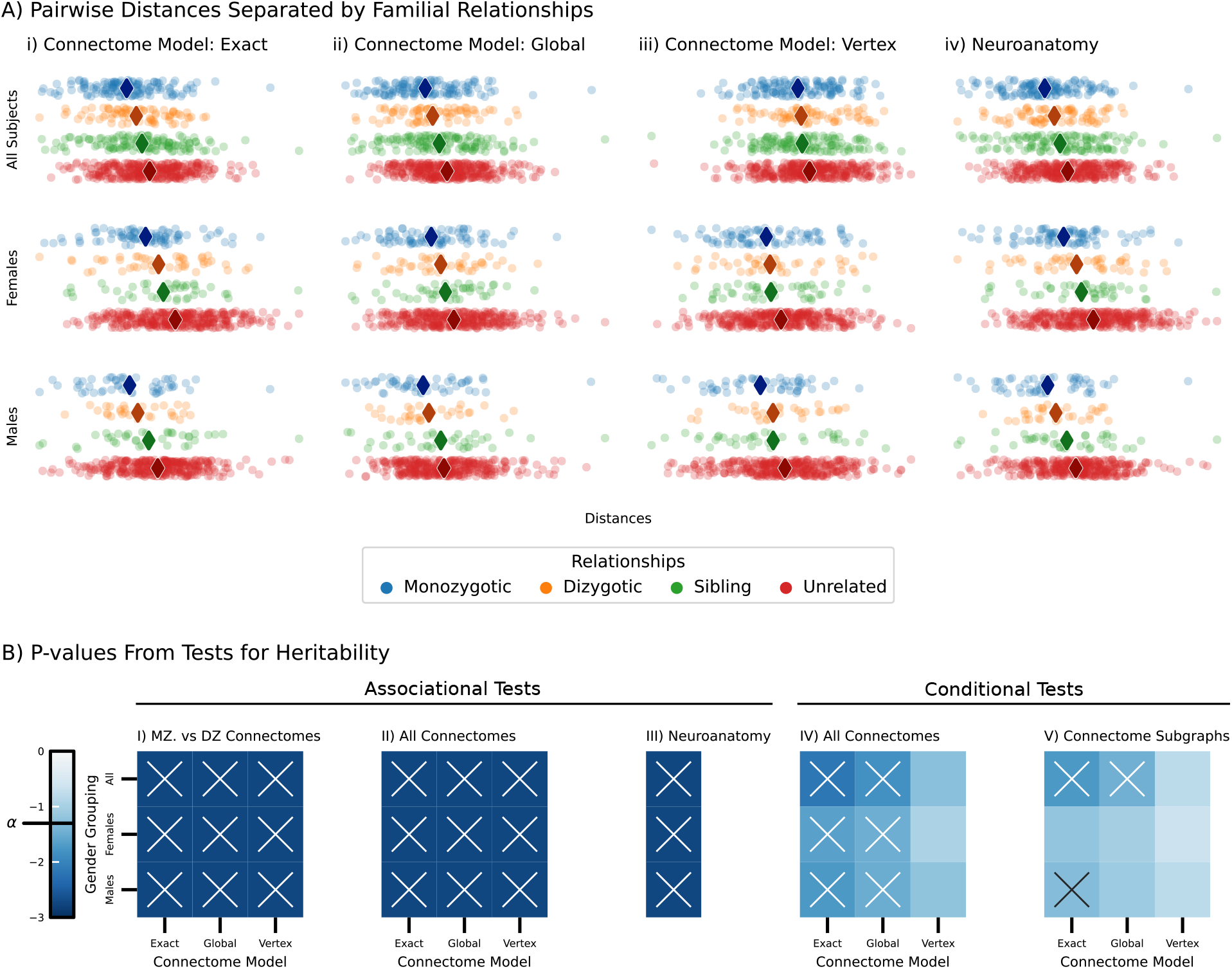
Testing associational and causal effect of genome on connectomes and neuroanatomy. **(A)** Each point represents the pairwise distance between pairs of participants; diamond markers represent the median distance, colors are familial relationships, and rows are sex. **(A.i-iii)** The median distances revealed that MZ twins exhibited smaller distances compared to DZ twins, siblings, and unrelated pairs, whereas the unrelated pairs showed larger distances. DZ twins displayed smaller distances compared to siblings and unrelated pairs, but larger than those of MZ twins. These findings provide qualitative evidence of heritability of connectomes. **(A.iv)** The median distances of neuroanatomy also show qualitative evidence of heritability. **(B)** Colors of heatmaps represent p-values, rows are gender groups, and columns are connectome models. (B.I-II) Associational test for connectome heritability. In (B.I), we only compare MZ and DZ twins, and the significance tests provide evidence of heritability as shown in (A.I-III). In (B.II), we conduct tests across all three familial relationships and observe significant effects, providing further evidence for connectomic heritability. In contrast, in (B.III), we observe significant effects for neuroanatomy, suggesting that the results in (B.I-II) may be confounded by an effect mediator. Subsequently, in (B.IV-V), we perform causal tests for connectome heritability. In (B.IV), we observe significant results only for the exact and global models, indicating that the connectomes remain heritable even after conditioning on the anatomical covariates, age, and sex. This suggests that the results in (B.I-II) cannot be entirely explained by neuroanatomy. The non-significant tests using the vertex model suggest that when we control for covariates and vertex-wise scaling, we can no longer detect heritability. In (B.V), we define a non-heritable subgraph as a collection of vertices that are not heritable by the causal test. We still observe several significant tests, suggesting heritability remains even after discarding vertices.

These results provide our first evidence for the heritability of connectomes. We then expanded our examination to include non-twin siblings and unrelated subjects (Figure 5A.I-III green and red rows) and observed that the medians grow larger as the familial relationships grow distant. Based on these observations, we tested associational heritability in all family relationships, producing p-values of *<* 10^*−*3^ for males and females under all three heritability models (Figure 5B.II). All of these tests indicate a strong associational effect of the genome on connectomes.

### 3.2 Associational Test for Neuroanatomic Heritability

Multiple studies have investigated the heritability of human brain anatomy [35–40]. Structural MRI studies on adults have shown that genetic and environmental factors have a significant impact on brain properties such as volume, shape, and both cortical and subcortical structures. This suggests that neuroanatomy could be an effect mediator (Figure 2). To assess if neuroanatomy could be a mediator, several features such as brain volume, axial diffusivity (AD), radial diffusivity (RD), and fractional anisotropy (FA) were computed for each region of the brain. Refer to Section 5.5 for more details.

We used Dcorr to test for the associational effect of the genome on anatomical covariates. This test assumes that if the covariates and genetics are independent, then brain anatomy is not heritable and has no effect on connectomes. Conversely, if the alternative is true, the associational effects of heritability of connectomes may be partially explained by the heritability of the anatomy itself, making the brain anatomy an effect mediator. Figure 5A.IV illustrates that similar to connectome distances, the median distances increase as familial relationships become more distant. Furthermore, we find that the tests showed *p <* 10^*−*3^ for all three gender groups under all three connectome models (Figure 5B.III).

The significance suggest that the associational heritability of connectomes may partly be due to the effect of anatomy on connectomes.

### 3.3 Causal test for connectomic heritability

To account for the dependence of neuroanatomy on genetics, we utilized conditional distance correlation (CDcorr) to estimate the conditional effect of genome on connectomes (Eq. 5.5). The covariate set also included age, which is a confounder. Our results showed *p <* 10^*−*2^ for exact and global models, but showed *p >* .1 for the vertex model. This indicates that while connectomes are heritable and not entirely influenced by anatomy and age, there is no significant underlying structure beyond the vertex-wise scaling. This contrasts with the results shown in Figure 5B.I-II, highlighting the importance of causal modeling in this study.

### 3.4 Non-heritable Subgraphs are also Heritable

In this section, we consider the possibility that heritability effects detected in Section 3.3 are driven by a subset of vertices rather than the entire connectome. That is, it is possible that the connectivity of a subset of brain regions may be highly similar in twins but highly different in non-twin siblings or unrelated individuals. To investigate this, we test for causal heritability by examining the *non-heritable subgraphs*, which are subgraphs induced by the vertices in which no heritable effect can be detected. See Section 5.1 for more details on the hypothesis.

The outcomes presented in Figure 5B.V display significant p-values for both the exact and global models across all genders, with only the exact model presenting significant p-values in males. These results suggest that for females, heritability can be fully accounted for by the heritable vertices, while in males, the non-heritable vertices still demonstrate heritability. Nevertheless, when controlling for scaling, the significance observed in the exact model for males is eliminated, indicating that differences in the scaling of the subgraphs explain the observed significance. These findings suggest the presence of an underlying structure in the non-heritable subgraph that cannot be detected at the level of individual vertices and highlights the necessity of statistical modeling of networks.

## 4. Discussion

### 4.1 Summary

In this study, we presented the first examination of the heritability of human connectomes using CHaiN. Our investigations into the heritability of connectomes provides strong evidence that the structural connectivity patterns within them are highly influenced by genetics by leveraging causal models and developing a test procedure to detect heritability. To establish this, we first compared the connectomes of monozygotic twins with same-sex dizygotic twins, which allowed us to account for shared environmental factors. Our results demonstrated that not only are connectomes reliable and meaningful, but they are also significantly influenced by genetics. We then extended our analysis to include monozygotic, dizygotic, and non-twin siblings, which further confirmed that connectomes are heritable. However, these results may be confounded by both observed and unobserved covariates. Therefore, we define a set of covariates, which we believe can account for all possible other unobserved covariates. Given this, we found that connectomes remained heritable even after controlling for age and the mediating effect of neuroanatomy. Finally, we demonstrated that the subgraphs formed by sets of brain regions that are not heritable within connectomes are also heritable. This suggests that there is an underlying structure within these subgraphs that is not evident in individual vertices.

A key innovation in this study is the utilization of statistical models, specifically the random dot product graphs, for connectomes, which introduces a new representation called latent positions. The models allow us to compare connectomes in various ways and explore different aspects of heritability. The simplest model we use is the exact model, which assesses any differences in the latent positions of connectomes. The global model, on the other hand, accounts for the possibility that connectomes may have different scales, such as variations in edge weights or more connections in general. Therefore, this model quantifies the differences in latent positions after removing the effect of global scaling. The vertex model allows for both global and vertex-wise scaling, enabling the three models to account for increasingly complex structures in connectomes as we move from the exact model to the vertex model. Our findings, as presented in Section 3.3, indicate that genetics play a crucial role in the differences of vertex scaling. However, when addressing vertex scaling of connectomes, heritability becomes undetectable, suggesting that heritability is only due to vertex scaling and not other underlying structures present in the connectomes. These results imply that previous conclusions on connectomic heritability may be influenced by vertex scaling and emphasize the need for further research into the relationship between vertex scaling in RDPG and network statistics, such as clustering coefficients and average path lengths.

There are several other open questions and opportunities for future research on the heritability of connectomes. One potential direction involves studying heritability in different neuroimaging modalities. Although our focus was on human connectomes estimated from diffusion MRI, the methods presented here can be applied to study heritability in connectomes obtained from other modalities, such as functional MRI, magnetoencephalography (MEG), and electroencephalography (EEG), or even in other species, like mice. Another potential direction is incorporating more detailed genetic data, including GWAS, to identify specific genetic variants associated with brain connectivity patterns. This would enable more direct and meaningful comparisons of genomes between subjects and could be readily integrated into the testing procedure presented here.

The findings on heritability in connectome analysis have significant implications for understanding the genetic basis of various neurological and psychiatric disorders. Disruptions in brain connectivity have been implicated in numerous conditions, such as autism spectrum disorder, schizophrenia, and Alzheimer’s disease [41, 42]. By revealing the heritable components of brain connectivity, researchers can start to pinpoint specific genetic factors that contribute to the development of these disorders. This knowledge could inform the creation of novel diagnostic tools, treatment strategies, and targeted interventions, ultimately enhancing the prognosis and quality of life for individuals affected by these disorders.

### 4.2 Limitations

#### Estimating Heritability

Heritability is typically defined as the relative contribution of genetic variations as opposed to environmental variations in phenotypic traits, and tests whether the heritability estimate is statistically significant, meaning if heritability exists or not. Numerous methods have been developed to partition phenotypic variation into genetic and environmental components, including techniques such as structural equation modeling (SEM). However, the causal approach presented in this study is distinct from these methods, as it does not aim to estimate heritability in a quantitative manner. Instead, our approach focuses on identifying the presence or absence of heritability by examining causal relationships between genetic factors and phenotypic traits.

The causal approach presented in this study offers a complementary perspective that can contribute to resolving the missing heritability problem. The missing heritability problem arises when the estimated heritability from genetic markers, such as those identified in genome-wide association studies (GWAS), accounts for only a small fraction of the heritability inferred from twin studies using SEMs or similar methods. This discrepancy between the observed and expected heritability is a major challenge in genetic research, as it limits our understanding of the genetic architecture underlying complex traits and diseases. By focusing on the presence or absence of causal relationships between genetic factors and phenotypic traits, this approach can help to identify previously unexplored genetic influences that might not be captured by traditional heritability estimation methods.

#### Choice of Model

We utilized the random dot product graph (RDPG) model in our analysis, which provided a unified framework for analyzing connectomes at both the network and individual vertex levels. However, it is important to acknowledge that every model has its limitations. The RDPG model can only represent a subset of all possible stochastic block models, and may not be able to capture certain patterns in connectivity. For example, if human brains were characterized by more connections between hemispheres than within hemispheres, the RDPG model will not be able to capture these patterns. To address these limitations, other models have been developed, such as the generalized random dot product graph, which can represent a broader range of stochastic block models and other network models. Additionally, there are models that are designed to analyze populations of networks, such as the common subspace independent edge model and joint random dot product graph model. Choosing any of these models enables one to define a valid distance metric for comparing connectomes. It is worth noting that there is currently no consensus on which model is best suited for modeling the human brain. Therefore, researchers must carefully consider the advantages and limitations of each model when selecting the appropriate one for their analysis. Ultimately, the choice of model will depend on the research question, the data available, and the assumptions that are deemed reasonable for the specific context.

## 5 Methods

### 5.1 Causal Analysis for Heritability

Throughout this paper, we use *P*_*X*_ to denote a generic distribution function and *f*_*X*_ to denote a generic probability density or mass function for a random variable *X*, which we abbreviate *P* and *f* for simplicity. We use *P*_*X*|*y*_ to denote the conditional distribution function of the random variable *X* conditioned on *Y* = *y*. The corresponding conditional density/mass function is *f* (*Y* = *y*|*X* = *x*), which we abbreviate *f* (*y*|*x*). We let *X* denote the *x* ∈ 𝒳 -valued random variable denoting the exposure, *Y* denote the *y* ∈ 𝒴-valued random variable denoting the outcome, *Z* denote the *z* ∈ *Ƶ*-valued random variable denoting the unmeasured covariates, and *W* denote the *w* ∈ *W*-valued random variable denoting the measured covariates. Throughout, we assume that (*X, Y, Z, W*) are sampled identically and independently from some true but unknown distribution (which implies exchangeability). In our investigation, the exposure *X* is the genome conferred to an individual, the outcome *Y* is an observed trait, such as the structural connectome, and *W* are covariates regarding the individual which are known (sex, neuroanatomy, and age). For more details on definitions, please see Bridgeford et al. [43].

We are interested in estimating the effect of different exposures on the outcome of interest, which can be quantified using the backdoor formula under the assumption that *W* and *Z* close all backdoor paths

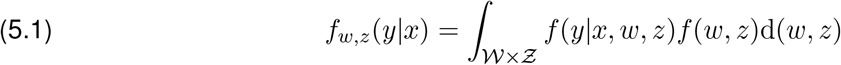

Note that the above equation integrates over the *unconditional* distribution of *W* and *Z, f* (*w, z*). This quantity can therefore be thought of as the distribution of the true outcome averaged over *all* measured and unmeasured covariates. This contrasts from the notationally similar, but practically distinct, quantity *f* (*y*|*x*), e.g.:

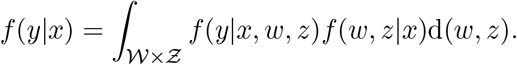

The key difference is that unlike Equation (5.1), *f* (*y*|*x*) averages the true outcome distribution, *f* (*y*|*x, w, z*), over the *conditional* distribution of the measured and unmeasured covariates, *f* (*w, z*|*x*). This has the implication that if *f* (*w, z*|*x*) is not the same as *f* (*w, z*|*x*′), *f* (*y*|*x*) represents an average over a different covariate distribution than *f* (*y*|*x*′).

#### Idealized Causal Heritability

Given the setup above, we say that *causal heritability* of a trait exists if and only if:

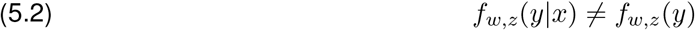

and we can construct a hypothesis as follows:

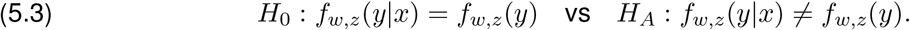

The null hypothesis states that there is no average causal effect; that is, the genome (exposure) has no effect on the connectomes, and this test can be formalized as an independence test. The key limitation of this characterization of heritability is that the estimand of interest *f*_*w,z*_(*y*|*x*) requires an integration over the unconditional distribution of the measured and unmeasured covariates, *f* (*w, z*), which we will not have in practice due to the fact that the covariates *Z* are unknown. Therefore, the above hypothesis test is not feasible since it would not be possible to formulate principled estimators of the estimand *f*_*w,z*_(*y*|*x*). In the following sections, we establish a series of related effects while establishing the specific conditions under which the existence of causal heritability effect can be inferred from the effect of interest. It is important to note that the validity of any causal assertions is contingent upon the extent to which these underlying assumptions accurately represent the data.

#### Associational Heritability

We only observe the pairs (*x*_*i*_, *y*_*i*_) for *i* ∈ [*n*]. Therefore we will only be able estimate functions of (*X, Y*). We say that *associational heritability* of a trait exists if:

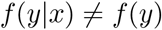

and we can construct a hypothesis as follows:

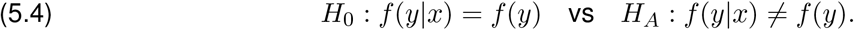

Under what circumstances can associational heritability be considered equal to causal heritability? A sufficient assumption for the equivalence of these two effects is that the measured and unmeasured covariates (*W, Z*) are non-confounding, which is the case when (*X, Y*) is independent of (*W, Z*). However, this assumption is often not reasonable. For example, if the size of human brains is associated with structural connectomes, then differences in *f* (*y*|*x*) and *f* (*y*) could be due to genetics or neuroanatomy, making it challenging to differentiate between causal and associational effects. This motivates the following approach.

#### Conditional Heritability

We observe the triples (*x*_*i*_, *y*_*i*_, *w*_*i*_) for *i* ∈ [*n*]. Thus, we will only be able to estimate the functions of (*X, Y, W*). Thus, a *conditional heritability* of a trait exists if:

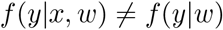

and we can construct a hypothesis as follows:

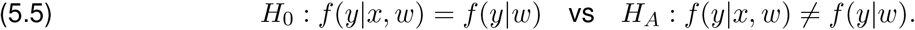

Under what circumstances can conditional heritability be considered equal to causal heritability? A sufficient condition for these two effects to be equivalent is that the strong ignorability assumption is satisfied [32]. Strong ignorability consists of two conditions:

1. The exposure is independent of the potential outcomes, conditioned on the observed and unobserved covariates (*W, Z*): *X ╨ Y* |*W, Z*.
2. The distributions of the observed and unobserved covariates have sufficient overlap among the exposure. Specifically, for all (*w, z*) ∈ *𝒳 × Ƶ*, 0 *<* ℙ(*X* = *x*|*W* = *w, Z* = *z*) *<* 1.

The first condition of strong ignorability can be specified in terms of backdoor paths, which are a set of paths between the exposure and the outcome that are not part of the causal pathway. These paths can create a spurious correlation between exposure and outcome if they are not properly controlled for. Therefore, if all backdoor paths between the exposure and outcome are blocked by conditioning on a set of observed covariates, then the exposure is rendered independent of potential outcomes. In other words, the observed covariates are assumed to be sufficient to control for all confounding variables, including unmeasured covariates, that could influence exposure and outcome. Thus, the main limitation of the conditional effect is that there must be no unmeasured confounding, which is an untestable assumption.

In the context of backdoor paths, the covariate overlap condition ensures that there is adequate variation in the covariates to block all the backdoor paths between the exposures and the outcomes. For example, if there is a subset of the population with extreme values of the covariates where no one is given a particular exposure, then it will be impossible to estimate the effect of the exposure in that subgroup.

#### Conditional Heritability for Vertices

Here We observe the triples (*x*_*i*_, *y*_*i*_, *w*_*i*_) for *i* ∈ [*n*], but instead consider each individual vertex, or a region of the brain. Thus, condiTo identify the non-heritable sub-graphs, we modify the hypothesis in Eq. 5.5 to allow for vertex-wise testing:

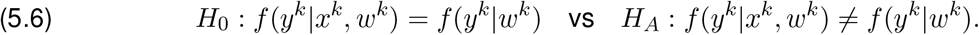

where *k* indexes a particular vertex, and *y*^*k*^ is a matrix of latent positions from all connectomes for vertex *k*. After computing a p-value for each vertex, those with p-values less than *α* = 0.05 after multiple hypothesis correction were discarded. The remaining set of vertices induces the non-heritable subgraphs. We then tested for conditional heritability of the non-heritable subgraphs as in Eq. 5.5.

#### Hypothesis Tests for Discovering Heritability

Classical statistical tests such as Student’s t-test, one-way analysis of variance (ANOVA), and their multivariate counterparts, such as Hotelling’s *T* ^2^ test and multivariate analysis of variance (MANOVA), are not well-suited for testing heritability in twin studies due to the inherent structure of the data. Specifically, the familial relationships (e.g. monozygotic twins, dizygotic twins) depend on which pair of subjects being examined rather than the individuals. For instance, consider two families, each with a set of monozygotic twins. Within families, the subjects are considered as monozygotic twins, but when compared across families, they are considered as unrelated subjects. Consequently, although both pairs are classified as monozygotic twins, they cannot be combined into a group for subsequent analysis. To address this issue, some studies examine within family pair differences rather than individual observations. However, this approach introduces another challenge: the pairwise differences are not independent. For example, in a family with one monozygotic twin pair and a non-twin sibling, there are three possible pairings (one twin pair and two non-twin sibling pairs). When comparing the connectomes of all possible pairs among the three individuals, the twins would be used for comparison in both the monozygotic and sibling groups.

Distance correlation (Dcorr) is a non-parametric test that can detect linear and non-linear dependency between two multivariate variables, and resolves the issue of pairwise dependence [44, 45]. The first step in Dcorr is to compute the distance matrices for the variables being compared. These matrices represent the distances, or differences, between all possible pairs of observations in the dataset. By working with distance matrices, Dcorr inherently takes into account the relationships between pairs of observations. Therefore, we used distance correlation to test the hypothesis of associational effect (Eq. 5.4). To test the hypothesis of causal effect (Eq. 5.5), we use the conditional distance correlation (CDcorr), which augments the Dcorr procedure by conditioning on the kernel of third variable [46]. In the following section, we present models and methods of computing distances between a pair of connectomes.

### 5.2 Network Construction

Connectomes were estimated using the ndmg pipeline [47], which is designed to process and generate connectomes from human dMRI and sMRI that minimizes batch effects across datasets. We note that higher b-values than 1000 were discarded prior to preprocessing. The dMRI scans were first denoised from Gibbs ringing using DiPy’s gibbs-removal, and then were pre-processed for eddy currents using FSL’s eddy-correct [48]. The results were denoised using DiPy’s Local PCA. FSL’s “standard” linear registration pipeline was used to register the sMRI and dMRI images to the MNI152 atlas [48–51]. A constant solid angle orientation distribution function (CSA-ODF) model is fit using DiPy [52] to obtain an estimated tensor at each voxel. A local deterministic tractography algorithm is applied using DiPy’s [52] to obtain streamlines, which indicate the voxels connected by an axonal fiber tract. Given a parcellation with vertices *V* and a corresponding mapping *P* (*v*_*i*_) indicating the voxels within a region *i*, we contract our fiber streamlines as follows. 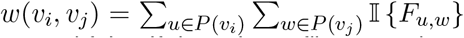 where *F*_*u,w*_ is true if a fiber tract exists between voxels *u* and *w*, and false if there is no fiber tract between voxels *u* and *w*. Several parcellations were used to generate connectomes, such as the Schaefer [53], AAL [54], Glasser [55] and a projection of the Desikan parcellation [56] to sub-cortical white-matter structures [57].

### 5.3 Network Preliminaries

Networks (or graphs) are convenient mathematical objects for representing connectomes. A network *G* consists of an ordered set (*V, E*) where *V* is the vertex set, and *E* is the set of edges. The set of vertices is represented as *V* = *{*1, 2, …, *n}* where |*V* | = *n*. The set of edges is a subset of all possible edges (e.g. *E ⊂ V × V*). An edge exists between vertex *i* and *j* if (*i, j*) ∈ *E*. A network can also be represented by its adjacency matrix **A** ∈ ℝ^*n×n*^ where each element, **A**_*ij*_, represents the edge between vertices *i* and *j*. In many connectomics datasets, edges have associated weights that represents some notion of strength of connections between two vertices. For example, edge weights from structural connectomes are non-negative integers that represent the number of estimated fiber tracts between two brain regions.

### 5.4 Statistical Model for Connectomes

Connectomes can be modeled using statistical models designed for network data. These statistical models consider the entire network as a random variable, including the inherent structure, dependencies within networks, and the noise in observed data [31, 58, 59]. In the following sections, we define the *random dot product graph* (RDPG) [60–62], how the parameters can be estimated from observed data, and then define the testing procedure.

#### Random Dot Product Graph Model

The (RDPG) is a type of independent edge model. In this model, each element of an adjacency matrix is sampled independently from a Bernoulli distribution:

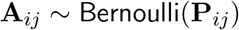

Given the number of vertices *n*, the matrix **P** is a *n × n* matrix of edge-wise connection probabilities with elements in [0, 1]. We can construct various models depending on the constraints imposed on **P**. Note that we assume that **P** has no self-loops (i.e. diag 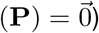 and is undirected (i.e. **P**^*T*^ = **P**).

In the random dot product graph (RDPG), the probability of a connection **P**_*ij*_ is determined by the vertices. Each vertex *i* ∈ *V* is associated with a low-dimensional *latent position* vector, **X**_*i*_, in the Euclidean space ℝ^*d*^. The probability of connection between vertices *i* and *j* is given by the dot product (i.e 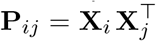). Thus, in a *d* – dimensional RDPG with *n* vertices, the rows of the matrix **X** ∈ ℝ^*n×n*^ are the latent positions of each vertex, and the matrix of edge-wise connection probabilities is given by **P** = **X X**^*T*^. Each element of the adjacency matrix is then independently modelled as

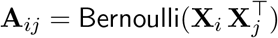

where **X**_*i*_ and **X**_*j*_ are latent positions for vertices *i* and *j*, respectively. We acknowledge that the original intention of RDPG is to model binary networks, although the model can be naturally extended to handle weighted networks. However, the weighted RDPG models are not well studied, and as such does not enjoy the same statistical guarantees. In the subsequent section, we present an algorithm for estimating latent positions from observed data and describe methods for preprocessing the data to enable interpretation of results within the context of binary networks while still utilizing weighted network data.

#### Adjacency Spectral Embedding

The modeling assumptions of the RDPG make the estimation of latent positions, which are usually unobserved in practice, analytically tractable. The estimation procedure we use is *adjacency spectral embedding* (ASE) [63]. The ASE of an adjacency matrix **A** into *d*-dimensions is 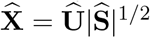 where 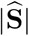 is a diagonal *d × d* matrix containing on the main diagonal the absolute value of the top-*d* eigenvalues of **A** in magnitude, in decreasing order, and Û is an *n × d* matrix containing the corresponding orthonormal eigenvectors. This simple and computationally efficient approach results in consistent estimates 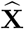 of the true latent positions **X** [63–65]. The ASE depends on a parameter *d* that corresponds to the rank of the expected adjacency matrix conditional on the latent positions; in practice, we estimate this dimension, 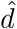, via the scree plot of the eigenvalues of the adjacency matrix which can be done automatically via a likelihood profile approach [66].

Structural connectomes have weighted edges (i.e. non-negative integers) and have no self-loops. To improve estimation of latent positions via ASE, we implement two preprocessing steps in order: 1) *pass-to-ranks* (PTR), and 2) *diagonal augmentation* (DA). PTR is a method for rescaling the *positive* edge weights such that all edge weights are in [0, 1]. The motivation of PTR is to convert the edge weights such that its distribution is uniform [47, 67]. Given an adjacency matrix **A** ∈ ℝ^*n×n*^, let *R*(**A**_*ij*_) be the “rank” of **A**_*ij*_, that is, *R*(**A**_*ij*_) = *k* if **A**_*ij*_ is the *k*^*th*^ smallest element in **A**. The rescaled adjacency matrix, **Ã** is:

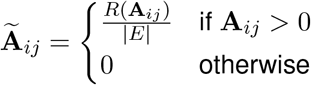

where |*E* | is the number of non-zero edges. Ties in rank are broken by averaging the ranks. DA is a method for augmenting the diagonal of an adjacency matrix. Given an adjacency matrix **A** ∈ ℝ^*n×n*^, the diagonal augmented matrix 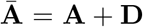 where **D** ∈ ℝ^*n×n*^ is a diagonal matrix with entries 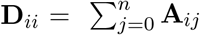

### 5.5 Distance Measures for Distance Correlation and Conditional Distance Correlation

The hypothesis testing procedures in Section 5.1 require functions to quantify dissimilarity (distances) or similarities between pairs of observations. In the following sections, we detail the functions used to compute the distance matrices for connectomes and genomes, and the kernel matrix for covariates.

#### Distances of Connectomes

In [68], the authors proposed three methods for calculating the distance between two networks under the `RDPG` model. Rather than comparing the high-dimensional connectomes directly, we first obtain low-dimensional latent positions of the connectomes via ASE and then compared these new representations. Each method makes different assumptions about the structure of the connectomes, providing different interpretations of the results.

Let **A**^(1)^ and **A**^(2)^ be *n × n* adjacency matrices, and let 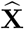 and 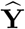 be *n × d* latent position matrices from their respective ASE. The distance metrics are defined as:

#### 1 Exact Model

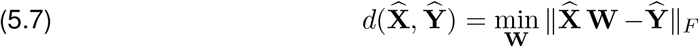

where **W** is a *d × d* orthogonal matrix that is estimated by solving the orthogonal Procrustes problem.

#### 2 Global Model

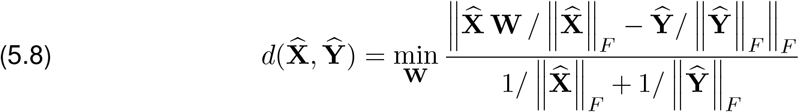

Note that a scaling constant is not estimated directly but the effect of scaling is removed by ensuring all latent position matrices have the same scale by normalizing by their respective Frobenius norms.

#### 3 Vertex Model

For any matrix **Z** with shape *n × d*, we define the 𝒟 (**Z**) as a diagonal *n × n* matrix whose main diagonal entries are the Euclidean norm of the rows of **Z**. Let 𝒟 ^*−*1^(**Z**) be defined as 1*/*(min_*i*_ ||**Z**_*i*_||), and let *𝒫* (**Z**) be the matrix whose rows are the projection of the rows of **Z** onto the unit sphere. Then the distance is defined as:

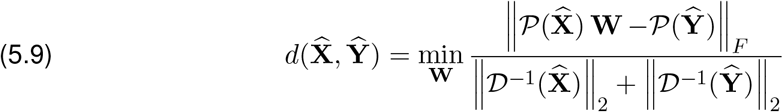

#### Distances of Genomes

Genomes are costly and challenging to measure, but their similarity can be estimated even without measurements by making strong assumptions. The coefficient of kinship (*φ*_*ij*_) is commonly used, which represents the probability that two individuals (subjects *i* and *j*) have the same allele or a DNA sequence at a random position in the genome [69]. For instance, monozygotic twins have an identical genome but two possible alleles since humans have two copies of a gene, resulting in a coefficient of *φ*_*ij*_ = 1*/*2. On the other hand, dizygotic twins and siblings share 50% of the genome on average, resulting in a coefficient of *φ*_*ij*_ = 1*/*4. Therefore, the distance between two genomes *G*_*i*_ and *G*_*j*_ is defined as:

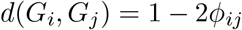

Table 1 enumerates all the possible distances. In the context of distance correlations, we note that any monotonic transformations of the values given to the genetic distance are equivalent as long as the ordinality remains.

**Table 1:** Genomic distances given by coefficient of kinships.

| Relationship | $\phi_{ij}$ | $1 - 2\phi_{ij}$ |
| --- | --- | --- |
| Monozygotic | $\frac{1}{2}$ | 0 |
| Dizygotic | $\frac{1}{4}$ | $\frac{1}{2}$ |
| Siblings | $\frac{1}{4}$ | $\frac{1}{2}$ |
| Unrelated | 0 | 1 |

#### Similarity/Distances of Covariates

In Section 3.2, we explored how genomes influence neuroanatomy by measuring brain volume, radial diffusivity (RD), axial diffusivity (AD), and fractional anisotropy (FA). These features were estimated per region of interest (ROI) defined by a parcellation, using the processed images obtained during network construction. RD is a measure of diffusion perpendicular to the axonal fibers, and it reflects the degree of damage or demyelination in the white matter. AD, on the other hand, is a measure of diffusion parallel to the axonal fibers and reflects the integrity and density of the axonal fibers. FA is a measure of the degree of anisotropy or directionality of water diffusion and reflects the coherence and organization of the white matter. We computed the average of these features across all regions, and we normalized the data by dividing each feature by its maximum value. In order to perform the hypothesis test using Dcorr in Section 3.2, we first compute the distance matrix for neuroanatomy. Let *w*_*i*_, *w*_*j*_ ∈ ℝ^*d*^ be measurements of observed covariates from subject *i* and *j* with dimensionality *d*. Then the distance function is simply given by the Euclidean norm:

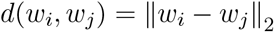

On the other hand, CDcorr requires a similarity matrix of the conditioning covariates, which includes the four neuroanatomy features and age. Age is normalized by dividing by the maximum age of the dataset. As before, let *w*_*i*_, *w*_*j*_ ∈ ℝ^*d*^ be measurements of observed covariates from subject *i* and *j* with dimensionality *d*. The kernel function is

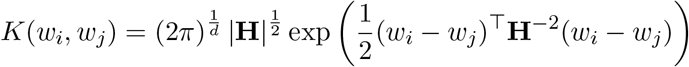

where **H** is a *d × d* diagonal matrix containing on the main diagonal the kernel bandwidth (*h*_1_, …, *h*_*d*_). While many methods exist for estimating the bandwidth, we chose a plug-in rule given by [70]:

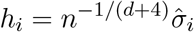

where *n* is the number of observations, *d* is the dimensionality of each observation, and 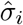 is the estimated standard deviation for dimension *i*.

### 5.6 *p* - values and Multiple Hypothesis Correction

All *p*-values from Dcorr and CDcorr tests are estimated using *N* = 25,000 permutations. Across all figures and tables associated with this work, we are concerned with obtaining a proper estimate of the rate at which we detect effects (*discoveries*).

Therefore, we control the false discovery rate (FDR) with the Benjamini-Hochberg Correction [71].

## 6 Data and Code Availability Statement

Analyses relied on graspologic [72], hyppo [73], NumPy [74], SciPy [75], Pandas [76], and NetworkX [77]. Plotting was performed using matplotlib [78] and Seaborn [79]. Processed connectomes can be found at s3://open-neurodata/hcp1200/. The data that support the findings of this study are available on request from the corresponding author JC. The data regarding subject-level data (e.g. age, family identifiers, sibling relationship identifiers) are restricted and not publicly available because the data might allow identification of individuals. All code used for this paper can be found at https://github.com/neurodata/connectomic-heritability and viewed as a JupyterBook [80] at http://docs.neurodata.io/connectomic-heritability.

## 7 Author Contributions

## 8 Acknowledgments

The authors are grateful for the support from the National Science Foundation (NSF) administered through NSF CAREER Award (Grant no. 1942963), the National Institute of Health (NIH) BRAIN Initiative (1RF1MH123233-01), and the National Institute of Health (NIH) through the National Institute of Mental Health (NIMH) Research Project RF1MH128696-01. Data were provided [in part] by the Human Connectome Project, WU-Minn Consortium (Principal Investigators: David Van Essen and Kamil Ugurbil; 1U54MH091657) funded by the 16 NIH Institutes and Centers that support the NIH Blueprint for Neuroscience Research; and by the McDonnell Center for Systems Neuroscience at Washington University.

## Appendix A. Simulated Connectomes

In this section, we describe the parameters that were used to generate the simulated connectomes in §2.3. In the following sections, we describe the formulation of an SBM as an RDPG and introduce the parameters used to generate simulated connectomes.

### A.1 Stochastic Block Model

*Stochastic block model* (SBM) is a popular statistical model for networks [81]. In this model, the connection probability between vertex *i* and *j* is solely determined by which community *i* and *j* belong to. For *K* communities, the vertices are divided into blocks, which is either deterministic or random, by the community membership indicators 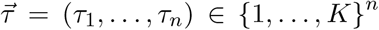 The matrix of probabilities is given by a *K × K* matrix **B** with entries in [0, 1]. Each element of the adjacency matrix is then independently modelled as

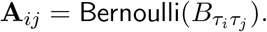

An SBM can be also be parameterized as an RDPG. In this case, community *l* ∈ [*K*] has a *d*-dimensional latent vector *x*_*l*_, and 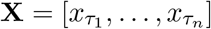 is an *n×d* matrix of latent positions where each row corresponds to the community latent position vertex *i*. We note that *d* is at most *K*. Therefore, the adjacency matrix is modelled as

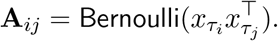

### A.2 Simulation Parameters

For all of the following simulations, we let the number of vertices *n* = 50 and the number of communities *K* = 2 with equal community sizes (e.g. 25 vertices per community). The communities in the simulated connectomes can be thought of as a hemisphere (e.g. left and right) of the brain, and each vertex is a region within one of the two hemispheres. For each set of simulation parameters, we generated 100 simulations. The average connectomes for simulated subjects 1 and 2 are shown in Figure 4(i) and (ii) columns. We then computed the difference in their estimated positions based on the three models of connectomes, which are also averaged. The resulting average differences are shown in Figure 4(iii-v) columns.

#### Exactly Same

The latent position for vertices belonging to community 1 is *x*_1_ = [1*/*4, 3*/*4] and those belonging to community 2 is *x*_2_ = [3*/*4, 1*/*4]. This results in the following block probability matrix:

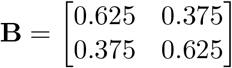

These parameters simulate a simplified brain; that is, the number of connections within each hemisphere is larger than across hemispheres [82]. If *U* and *V* denote adjacency matrices of subjects 1 and 2, then

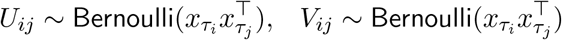

where *τ*_*i*_ represents the community assignment for vertex *i*. In this case, the parameters of the two subjects are exactly the same.

#### Same up to Global Scaling

Here, we introduce a global scale constant *c >* 0. The constant scales the connection probabilities of one subject. Then the adjacency matrices are sampled as:

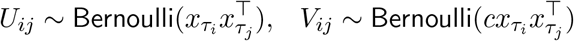

with the same latent position vectors as above. We set *c* = 1.2. Therefore, the two subjects have the same probability matrix conditioned on *c*.

#### Same up to Vertex Scaling

Here, we introduce a degree-correction parameter that controls the expected degree of a vertex (e.g. number of edges that connect to a vertex) *θ*_*i*_ ∈ [0, 1]. This model is known as the degree-corrected stochastic block model (DCSBM). The degree correction parameters are defined as 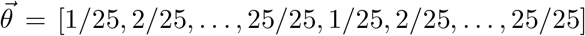 forming a vector with length *n*. Given the degree corrections, the adjacency matrices are sampled as:

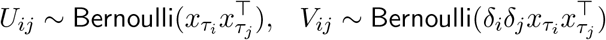

Therefore, the two subjects have the same probability matrix conditioned on 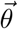

#### Parameters for Different Connectomes

Here we introduce a different set of latent position for one of the subjects: *y*_1_ = [4*/*5, 2*/*5] and *y*_2_ = [2*/*5, 2*/*5]. Given the same degree correction *θ*_*i*_ and *x*_1_, *x*_2_ as above, the adjacency matrices are sampled as:

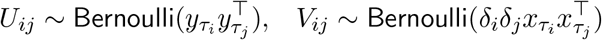

Therefore, the two subjects do not have the same probability matrix even after controlling for degree correction.

